# Motor unit discharge rate modulation during isometric contractions to failure is intensity and task dependent

**DOI:** 10.1101/2023.12.04.569929

**Authors:** Tamara Valenčič, Paul Ansdell, Callum G Brownstein, Padraig M Spillane, Aleš Holobar, Jakob Škarabot

## Abstract

The nature of neuromuscular decrements associated with contractions to task failure is known to dependent on task demands. Task-specificity of the associated adjustments in motor unit discharge rate (MUDR) behaviour, however, remains unclear. This study examined MUDR adjustments during different submaximal isometric knee-extension tasks to failure. Participants performed a sustained and an intermittent task at 20 and 50% of maximal voluntary torque (MVT), respectively (Experiment 1). High-density surface electromyography signals were recorded from vastus lateralis (VL) and medialis (VM) and decomposed into individual MU discharge timings, with the identified MUs tracked from recruitment to task failure. MUDR was quantified and normalised to intervals of 10% of contraction time (CT). MUDR of both muscles exhibited distinct modulation patterns in each task. During 20% MVT sustained task, MUDR decreased until ∼50% CT, after which it gradually returned to baseline. Conversely, during the 50% MVT intermittent task, MUDR remained stable until ∼40-50% CT, after which it started to continually increase until task failure. To explore the effect of contraction intensity on the observed patterns, VL and VM MUDR was quantified during sustained contractions at 30 and 50% MVT (Experiment 2). During the 30% MVT sustained task, MUDR remained stable until ∼80-90% CT in both muscles, after which it continually increased until task failure. During the 50% MVT sustained task the increase in MUDR occurred earlier, after ∼70-80% CT. Our results suggest that adjustments in MUDR during submaximal isometric contractions to failure are task- and intensity-dependent.

## INTRODUCTION

Voluntary muscle force or torque production is regulated via activity of spinal motoneurons and groups of muscle fibres innervated by individual axons (i.e., motor units). When recruited, motor units (MUs) modulate their discharge rate in response to excitatory and inhibitory synaptic inputs to the motoneuron pool from descending supraspinal pathways, spinal interneurons, and muscle afferents (Heckman & Enoka, 2012). With repeated activation, such as during prolonged muscle contractions, neural and contractile properties of the motor pathway exhibit alterations, leading to a progressive reduction in muscle force production capacity and, ultimately, task failure (Enoka & Stuart, 1992; Enoka & Duchateau, 2016). When motor output is submaximal, the net torque can be maintained by recruitment of additional MUs, and adjustments in their discharge rate via increases in the excitatory synaptic input (Enoka & Stuart, 1992; Hunter, 2018). Whilst previous literature unanimously reports progressive increases in the number of active MUs during prolonged sustained or intermittent submaximal isometric contractions, the reported patterns of MU discharge rate modulation during prolonged contractions have been inconsistent. Most commonly a gradual decrease (Garland *et al*., 1994; Carpentier *et al*., 2001; Vila-Chã *et al*., 2012; McManus *et al*., 2016; Lowe *et al*., 2023), increase (Bigland-Ritchie *et al*., 1986; Kuchinad *et al*., 2004), or a biphasic modulation pattern, i.e. an initial decrease followed by a progressive increase until task failure (Griffin *et al*., 2000; Adam & De Luca, 2005; Mettler & Griffin, 2016; Martinez-Valdes *et al*., 2020a) have been demonstrated. Considering the nature of neuromuscular decrements associated with contractions to task failure is dependent on task demands (Enoka & Stuart, 1992; Hunter, 2018), the differences in submaximal contraction intensities and characteristics between studies (i.e., task-dependency) might explain inconsistency of MU discharge behaviour reported in previous studies, but this remains to be tested directly.

The physiological responses to prolonged sustained and intermittent muscle contractions, even when performed in the same exercise intensity domain, differ due to differences in muscle perfusion and neural activation, with potential implications for MU discharge rate characteristics (Taylor *et al*., 2016; Hunter, 2018). Whilst prolonged sustained contractions critically limit muscle perfusion even at relatively low forces, causing muscle ischaemia and accelerated accumulation of contractile function-impairing anaerobic metabolites in the muscle, brief intermittent contractions are each followed by a hyperaemic response, which decelerates metabolic and ionic perturbations (Monod & Scherrer, 1965; Chidnok *et al*., 2013; Broxterman *et al*., 2014). Furthermore, differences in spinal reflex regulation and/or motoneuron activation between intermittent and sustained contraction could also influence MU discharge rate. For example, during sustained isometric contractions, motoneurons are tonically active, leading to intrinsically mediated reductions in responsiveness of motoneurons to constant synaptic input, shown as reduction in motoneuron discharge rate (i.e., spike frequency adaptation). Such an observation has been noted in both animal preparations using intra- or extracellular current injections (Kernell & Monster, 1982; Spielmann *et al*., 1993; Sawczuk *et al*., 1995; Button *et al*., 2007) and in humans as reductions in the size of evoked response to direct stimulation of corticospinal axons (Finn *et al*., 2018; Brownstein *et al*., 2021). Conversely, during intermittent contractions, motoneuron activation is interrupted by periods of rest. Using extracellular stimulation of spinal motoneurons in an anaesthetised cat preparation, intermittent stimulation has been demonstrated to lead to lower levels of spike frequency adaptation compared to constant stimulation (Spielmann *et al*., 1993). Furthermore, the greater metabolic demands of sustained contractions likely increase group III/IV afferent discharge, leading to greater reductions in motoneuron and cortical excitability (Martin *et al*., 2006; Hilty *et al*., 2011), and thus possibly affecting MU discharge rate modulation in a task specific manner. Indeed, greater muscle perfusion has been shown to induce smaller MU discharge rate modulation and lower MU discharge rate variability (Lowe *et al*., 2023). Similarly, excitatory spinal reflex input appears unaffected by intermittent contractions, but reduced during sustained contractions to failure (Duchateau *et al*., 2002), with a likely influence on task-specific MU discharge rate modulation patterns.

In this study, we aimed to examine and compare MU discharge rate modulation of the vastii muscles during sustained and intermittent submaximal isometric knee extension tasks to failure. To assess task-specificity of MU discharge rate adjustments (Experiment 1), the intensity of the tasks (20% and 50% of maximal voluntary torque [MVT] for sustained and intermittent task, respectively) was chosen to induce similar declines in MVT, voluntary activation, and muscle contractile properties at task failure (i.e., above the critical torque for the respective contraction modality, (Monod & Scherrer, 1965; Burnley, 2009). Because we could not exclude the possible confounding effects of contraction intensity to the observed MU discharge behaviour, we performed an additional experiment consisting of two sustained tasks to failure performed at different intensity levels (Experiment 2). For both experiments it was hypothesised that the time-dependent changes in MU discharge behaviour will differ between the tasks.

## MATERIALS AND METHODS

### Participants and ethical approval

Twenty-two healthy volunteers (11 males and 11 females) participated in Experiment 1. In thirteen participants, we were unable to identify sufficient number of MUs from the recorded signals (i.e., we were not able to track at least 1 MU throughout the entire task in at least one muscle), and thus the final sample on which results are reported here included nine participants (1 female; 26 (6) years, 1.75 (0.06) m, 73.9 (9.3) kg). In Experiment 2, eight healthy adults participated (1 female; 29 (4) years, 1.77 (0.08) m, 75.8 (12.0) kg).

Exclusion criteria for both experiments included any current injury to the lower limbs, back or spine, a history of a major traumatic injury or surgery, musculoskeletal or neuromuscular disorder affecting function of the major joints of the lower limb, and taking any medication known to influence neuromuscular function. To minimise the effects of menstrual cycle on neuromuscular function (Ansdell *et al*., 2019b; Piasecki *et al*., 2023), female participants were tested in the early follicular phase of the cycle (days 1-7) or during the period of contraceptive monophasic pill consumption. Before taking part in experimental procedures, participants provided written informed consent. The experiments were approved by Loughborough University Ethical Advisory Committee (2020-1733-1674 and 2023-14760-15109) and were conducted in accordance with the Declaration of Helsinki, except for registration in database.

### Experimental design and protocol

#### Experiment 1

Participants arrived to the laboratory on three separate occasions at a consistent time of day (± 1 hour), 2-10 days apart. Participants were instructed to arrive rested and hydrated, having avoided strenuous exercise, and refrained from alcohol and caffeine consumption for 24 and 12 hours prior to arrival, respectively. All sessions involved voluntary and evoked isometric knee extensions of the dominant limb (as determined by the lateral preference inventory (Coren, 1993)), with participants seated in a rigid, custom-built isometric dynamometer. The knee- and hip-joint angles were fixed at 115° and 126° (180° = full extension), respectively, with the testing chair adjusted to each participant’s stature and kept consistent across the sessions. During the performance of voluntary and evoked isometric contractions, participants were firmly strapped across the waist and chest to minimise extraneous movement.

The first visit was a familiarisation during which participants were habituated to percutaneous femoral nerve stimulation and practiced the performance of maximal and submaximal isometric knee extensions. Participants were also familiarised with the contraction tasks and therefore practiced the sustained contractions for a limited time; 30 and 120 s of the sustained (20% of maximal voluntary contraction, MVC) and intermittent (50% MVC) contraction tasks, respectively.

During the experimental sessions (visits 2 and 3), participants performed the contraction tasks until failure in a randomised, crossover design. Following a standardised warm-up of 3-second submaximal isometric knee extensions at 50% (×3), 75% (×3) and 90% (×1) of perceived maximal effort (separated by 15-30 seconds of rest), participants performed five ∼3-second MVCs separated by ≥30 seconds of rest to assess maximal voluntary force. During and ∼2 seconds after the final three MVCs, an electrical pulse was delivered to the femoral nerve to assess quadriceps voluntary activation level and evoked contractile muscle properties. To facilitate maximal force production, biofeedback was provided by displaying a real-time force trajectory on a computer monitor ∼1 m in front of the participant, with a horizontal cursor indicating the greatest force achieved. Participants were instructed to “push as hard as possible” and strong verbal encouragement was provided. The peak instantaneous force attained during the MVCs was used as a reference to calculate the submaximal force targets.

After a 5-minute rest period, participants performed either a sustained or an intermittent isometric knee extension task to failure at 50% and 20% MVC, respectively (randomised). These two specific contraction intensities were chosen to ensure participants performed contractions above the respective critical torque of the modality. It has been demonstrated that a critical torque corresponds to ∼15% MVC for sustained contractions (Monod & Scherrer, 1965), and ∼35% MVC for intermittent contractions (Burnley *et al*., 2012). Consequently, the intensities used in the present study were approximately 30-40% above previously reported critical torque values, with the aim of inducing similar reductions in maximal voluntary torque, voluntary activation, and potentiated twitch amplitude in both tasks, thus allowing the assessment of the modulation of MU discharge properties as a function of contraction task without the confounding effect of differences in decrements of neuromuscular function. In short, had we compared a sustained versus intermittent contractions at 20% MVC, the physiological demands and magnitude of neuromuscular impairments during the tasks would have differed, as 20% MVC intermittent contractions results in a target level below critical torque for most individuals (Burnley *et al*., 2012).

The intermittent contraction task involved a 2.75:1 (15-second) duty cycle consisting of a 3-second ramp increase in force output to the target level, a 5-second hold phase at the target level, and a 3-second ramp decrease to relaxation, followed by a 4-second rest. Participants were instructed to match a trapezoidal waveform with a real-time force trajectory displayed on the computer monitor. During the sustained contraction tasks, the force target was indicated by a horizontal line, which the participants were instructed to match as closely as possible. Time to task failure was recorded to the nearest second after three consecutive failures to meet the target force for the entire 5 seconds during the intermittent contraction task, and after a sustained 3-second decline in the force output by >10% of the target in the sustained contraction task, despite strong verbal encouragement. No feedback of the elapsed time was provided. Immediately after reaching task failure, participants performed three additional MVCs with percutaneous femoral nerve stimulation ∼10 seconds apart. HDsEMG recordings were performed throughout the sessions.

#### Experiment 2

Because the sustained and intermittent isometric contractions to failure in Experiment 1 were performed at different relative force levels, we could not exclude the potential influence of contraction level on the differential MU discharge modulation (see Results). Additionally, considering the decomposition bias towards identification of higher-threshold MUs (Francic & Holobar, 2021), we could not exclude differences in MU recruitment threshold as a factor in the observed MU discharge behaviour. Therefore, we performed an additional experiment where we assessed MU discharge behaviour during two isometric sustained contraction tasks at different contraction levels to failure. The main difference in Experiment 2 was the contraction intensity; here, participants performed sustained isometric knee extension to task failure at 30 and 50% MVC in a randomised, crossover design on two separate days (at least 48 hours, but no more than 10 days apart). Additionally, prior to these contraction tasks, and after the warm-up, participants performed two trapezoidal ramp contractions at 50 and 70% MVC, increasing the knee extensor force level at 10% MVC/s, maintaining the force level at the target for 10 s, and decreasing the force level at 10% MVC/s to resting levels. This allowed quantification of potential increases in MU discharge rate at task failure relative to MU discharge rate during a higher-intensity contraction in a fresh state. HDsEMG signals were recorded throughout the contraction tasks, and femoral nerve stimulation was delivered during and a few seconds after each of the three maximal effort contractions before and after the sustained isometric knee extension tasks.

### Experimental procedures

#### Force recordings

The force signal was measured using a calibrated S-beam strain gauge (linear range: 0-1.5 kN; Force Logic, Swallowfield, UK) attached perpendicularly to the participant’s tibia using a bespoke reinforced non-extendable webbing strap (35 mm width) tightly fastened superior to the lateral malleolus at ∼15% of the tibial length (distance between the lateral malleolus and the knee-joint centre). The analogue force signal was amplified (×370) and sampled at 2048 Hz simultaneously via an analogue-to-digital converter (Micro 1401; Cambridge Electronics Design Ltd., Cambridge, UK) for recordings of maximal voluntary force, voluntary activation and contractile twitch properties in Spike2 software (version 10; CED Ltd., Cambridge, UK), as well as via a 16-bit multichannel amplifier (Quattrocento, OT Bioelettronica, Torino, Italy) for synchronisation with HDsEMG signals.

#### Femoral nerve stimulation

Percutaneous electrical stimulation (single square-wave pulse, 1 ms, 300 V; DS7AH, Digitimer, Welwyn Garden City, UK) of the femoral nerve was delivered via circular self-adhesive gel-coated cathode and anode (32 mm^2^; CF3200, Nidd Valley Medical, Bordon, Hampshire, UK) positioned in the femoral triangle and mid-way between the greater trochanter and iliac crest, respectively. The stimulation current was determined by increasing the pulse intensity from 20 mA in 20-mA increments until a plateau in the resting quadriceps twitch force was attained. Stimulus intensity was then increased by 30% to ensure supramaximal stimulation.

#### High-density surface electromyography

The HDsEMG signals were recorded from the VL and VM muscles in monopolar configuration using semi-disposable 64-electrode grids (5 rows x 13 columns; 1 mm electrode diameter; 8 mm interelectrode distance; OT Bioelettronica, Torino, Italy) attached to the skin surface using disposable adhesive foam interfaces (SpesMedica, Battipaglia, Italy). Following skin preparation by shaving, light abrasion, and cleaning with ethanol, the adhesive foam interfaces were filled with conductive paste (SpesMedica), and the grids were positioned over the VL and VM muscles at ∼20° and ∼50° relative to the line between superior iliac spine and the lateral and medial border of the patella, respectively. The grid alignment with the orientation of fibres was confirmed by ultrasound (EUB-8500; Hitachi Medical Systems UK Ltd., Northamptonshire, UK; 92-mm linear-array transducer, EUP, L53L). Two self-adhesive reference electrodes (36 mm^2^; Cardinal Health, Dublin, OH, US) were attached across the patella of both limbs and the signal was grounded via a water-soaked strap electrode placed superior to the malleolus of the non-dominant limb. The HDsEMG signals were sampled at 2048 Hz and amplified using a 16-bit multichannel amplifier (Quattrocento, OT Bioelettronica, Torino, Italy; 3 dB), band-pass filtered (10-500 Hz), and recorded in OTBioLab+ software (OT Bioelettronica).

### Data analysis

#### Voluntary and evoked torque

Force data were low-pass filtered at 20 Hz (zero-lag, fourth-order Butterworth digital filter), gravity corrected, and converted to torque with multiplication by lever length (distance between the knee-joint centre and the centre of the ankle strap). The peak torque value achieved during maximal voluntary contractions was denoted as MVT. Voluntary activation level was quantified by comparing the amplitude of the superimposed twitch (SIT) with respect to the potentiated quadriceps twitch (Q_tw_) using the established equation (1 – [SIT / Q_tw_]) · 100 (Merton, 1954). A correction equation implementing torque at SIT (T_SIT_) was employed if the timing of the delivered stimulus did not correspond to the proximity of peak torque (100 – SIT · [T_SIT_ / MVT] / Q_tw_ · 100 (Strojnik & Komi, 1998). For MVT, VA and Q_tw_, the values of the three contractions before and after the fatiguing tasks, respectively, were averaged; unless a trial deviated from the average by more than 2 standard deviations in which case it was excluded from the calculation of mean. Torque steadiness during the two tasks was quantified as a coefficient of variation of torque (the ratio of standard deviation and mean torque levels) in the same time windows used for quantifying discharge rate characteristics (see below).

#### High-density surface EMG decomposition and motor unit tracking

Offline analyses were performed using MATLAB (R2023a; Mathworks Inc., MA). Monopolar HDsEMG signals were digitally band-pass filtered with a fourth-order, zero-lag Butterworth filter (20-500 Hz). We then removed channels exhibiting poor signal-to-noise ratio, noise, or artefacts using a semiautomated tool in MATLAB based on area under the power spectrum and amplitude; typically, this involved the removal of fewer than 5% of channels. HDsEMG signals were decomposed using the extensively validated Convolution Kernel Compensation algorithm (Holobar & Zazula, 2007). This algorithm is based on the blind source separation principles whereby the EMG mixing model is inverted, and MU filters are estimated, yielding the estimation of a MU spike train (Holobar & Farina, 2021; Škarabot *et al*., 2023a).

The calculation of MU filters and their application to a different portion of the signal from the same contraction or another contraction allows identification of discharges from the same MUs (Francic & Holobar, 2021), even in the case of different levels of MU synchronisation (Škarabot *et al*., 2023a). Thus, to track motor units across the contraction tasks to failure we used an approach of estimation and application of MU filters in shorter, overlapping windows (Rossato *et al*., 2022). For the sustained task, decomposition of the HDsEMG signal was performed independently in 30-second windows at the beginning and end of the signal (Figure 1A). From these decomposed sections of 30 seconds, MU spike trains were first visually inspected (Figure 1Bi), and MU filters iteratively optimised using standard procedures, segmenting the MU firings from noise/crosstalk from other MUs (Del Vecchio *et al*., 2020). After that, MU filters were iteratively applied in 15-second windows using a 50% overlap (Figure 1Biii) to the rest of the signal upon which the MU filters were again optimised before further application (Figure 1Biv).

**Figure 1.**
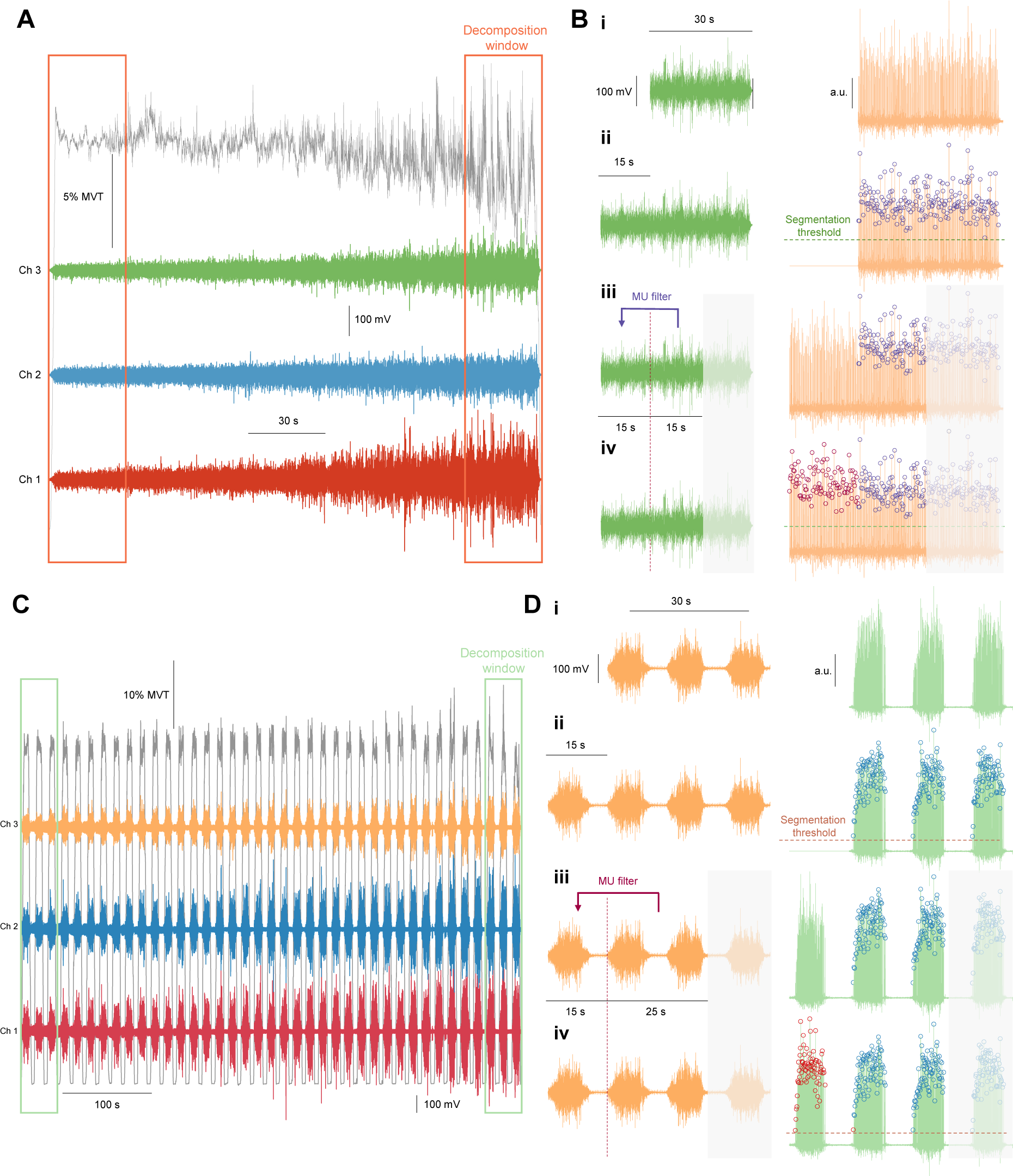
High-density electromyography decomposition and motor unit tracking. A, C: High-density electromyography (EMG) signals were decomposed in the 30-second or 41-second windows at the beginning and end of the sustained contraction task at 20% and the intermittent contraction task at 50% of maximal voluntary torque (MVT), respectively. For clarity, only 3 out of 64 EMG channels recorded from vastus lateralis and medialis for the sustained and intermittent tasks, respectively are shown in the example. B, D: The decomposed EMG signals yielded motor unit spike trains (i), which were then segmented into the identified motor unit discharges (denoted by circles in the right panel) and noise (ii), after which an overlapping window was used to apply the motor unit filters to the undecomposed portion of the EMG signal (iii), followed by another segmentation (iv). This process was iteratively completed for the remaining portion of the signal. Motor unit spike trains duplicates were identified in this process, followed by a removal of a duplicate motor unit with the lower pulse-to-noise ratio.

A similar approach was used for the intermittent task, however, here, 41-second portions of the signal (3 contractions; Figure 1C) were decomposed at the beginning and end of the task. MU spike trains were then visually inspected (Figure 1Di), with MU filters optimised through segmentation (Figure 1Dii), and then applied in 15-second windows (1 contraction) using a 30-second (2 contractions) overlap (Figure 1Diii), followed by another optimisation of MU filters (Figure 1Div). Because of successive independent decompositions (i.e., decomposition of the first and final portion of the tasks), several MU duplicates were inevitably identified. Duplicates were considered those that shared at least 30% of the same discharges (discharge match tolerance: 0.5 ms). Duplicates with the lower accuracy of MU spike train estimation (based on pulse-to-noise ratio; (Holobar *et al*., 2014) were discarded. Only MU spike train with a reliable discharge pattern and a pulse-to-noise ratio >28 dB were kept for analyses.

#### Discharge rate characteristics

Motor unit discharge rate was calculated as the mean of the reciprocal of the interspike interval in 1-second epochs during the stable region of the sustained and intermittent tasks (Figure 2). For the latter, this meant that the discharge rate was calculated in 1-second epochs during the plateau regions of the repeated trapezoidal contractions. Discharges involving interspike intervals >500 ms or <20 ms were excluded from analysis to minimise the confounding effect of sporadic discharges and/or potential decomposition errors on the subsequent calculations. Recruitment time was calculated as the time in the task at which a given MU started discharging (Figure 2). During the intermittent tasks, we could also reliably estimate recruitment and derecruitment thresholds, which were calculated as the mean torque level corresponding to the first and last three MU spikes, respectively. All variables were averaged using time-normalised intervals with length equal to 10% of total contraction time (i.e., 10-100% of total contraction time in 10% increments). For trapezoidal ramps performed before sustained tasks (Experiment 2), the average discharge rate during the 10-second plateau region of the contraction was calculated. For comparative purposes, similar calculation was performed on the final 10 seconds of the sustained tasks. For this analysis, we only compared discharge rate in the final 10 seconds of the sustained tasks with discharge rate during a trapezoidal ramp contraction at a level 20% greater than that of a sustained task (i.e., for 30% MVT sustained task comparison was made with a 50% MVT trapezoidal ramp contraction, whereas the 50% MVT sustained task was compared to a 70% MVT trapezoidal ramp).

**Figure 2.**
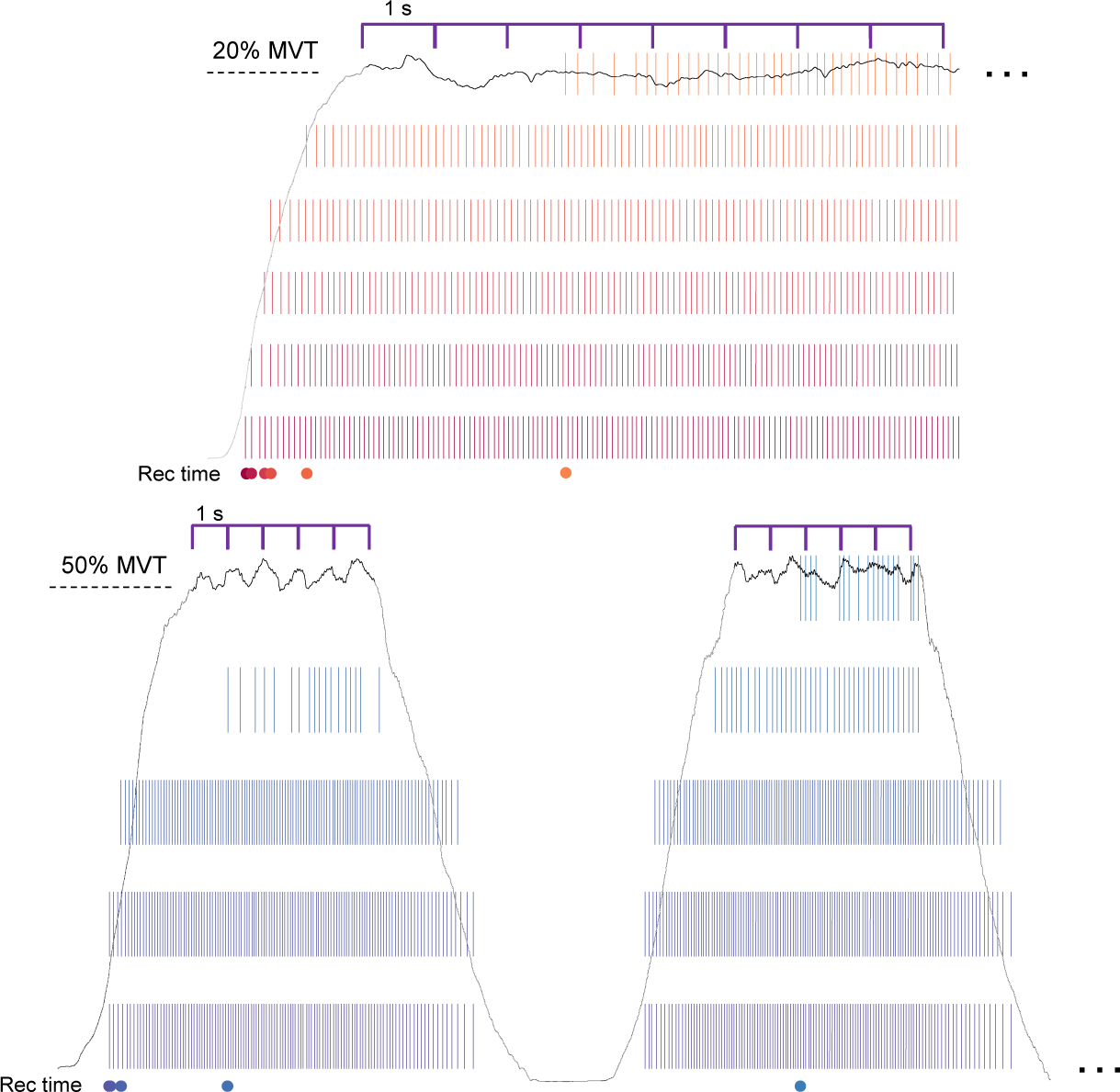
Analysis of motor unit discharge properties. Motor unit discharge rate was calculated as the mean of the reciprocal of the interspike interval in 1-second non-overlapping windows during sustained contraction tasks, and during the plateau regions of the intermittent task. Coefficient of variation of torque and root mean square electromyography signal amplitude were calculated in the same time windows. Recruitment time was denoted as the contraction time corresponding to the first spike of a motor unit.

#### Interference EMG amplitude

To characterise the interference EMG amplitude, we computed root mean square of a single bipolar recording derived from the HDsEMG signals. For this purpose, we averaged the activity of two sets of six channels in the central portion of the bidimensional electrode grid (columns 2 – 4, rows 4 – 5 and 6 – 7), then differentiated the signals (interelectrode distance 16 mm). Root mean square values were then calculated in the same time windows used for quantifying discharge rate characteristics (Figure 2). The interference EMG amplitude during the two exercise tasks was normalised to maximal interference EMG amplitude in a moving 1-second root mean square window during the performance of MVC.

### Statistical analysis

All statistical analyses were performed in R (R Foundation for Statistical Computing, Vienna, Austria). To assess differences in time to task failure, a paired t-test was performed. We used linear mixed effects models to assess whether discharge rate and interference EMG amplitude are predicted by fixed effects of contraction time (10-100%), task (intermittent, sustained), muscle (VL, VM) and their interaction, with participant used as a random intercept. Because additional MUs were recruited throughout the duration of the task, recruitment time was employed as a covariate. Modulation in recruitment and derecruitment thresholds was assessed for the intermittent task only, employing contraction time, muscle, and their interaction as fixed effects in the linear mixed model. Muscle was removed as a fixed factor when comparing modulation of torque steadiness during the two tasks. Changes in MVT, VA, and Q_tw_ were assessed using time (pre, post task), task, and their interaction as fixed effects. In Experiment 2, contraction level was used as a fixed factor instead of task. To compare differences in discharge rate during trapezoidal ramp contractions and the final 10 seconds of the sustained tasks, contraction (trapezoidal ramp, sustained), task (30% vs 50% MVT sustained), and muscle (VL, VM), and their interactions were considered as fixed effects, with participant taken as a random intercept. Significance of all models was assessed using analysis of variance with Satterthwaite’s method (*lmerTest* package; (Kuznetsova *et al*., 2017). The significance level was set at an alpha level of 0.05. In cases of significant main effects or interactions, pairwise post hoc tests of estimated marginal means with Tukey adjustment for multiple comparisons were performed (*emmeans* package; Lenth & Lenth, 2018).

## RESULTS

### Experiment 1

#### Time to task failure and decrements in neuromuscular function

Example torque traces of the intermittent and sustained tasks are depicted in Figure 3A. Time to task failure was 531 (134) and 217 (82) seconds for the intermittent and sustained task, respectively (t_8_ = −7.0, p = 0.0001). The knee extensor MVT (F_1, 24_ = 180.0, p < 0.0001; Figure 3B), voluntary activation level (F_1, 21_ = 22.8, p = 0.0001; Figure 3C), and potentiated twitch amplitude (F_1, 21_ = 32.0, p < 0.0001; Figure 3D) decreased after fatiguing tasks. No significant interactions were found between task and time to task failure (p ≥ 0.1516), suggesting the tasks induced similar levels of neuromuscular function decrements.

**Figure 3.**
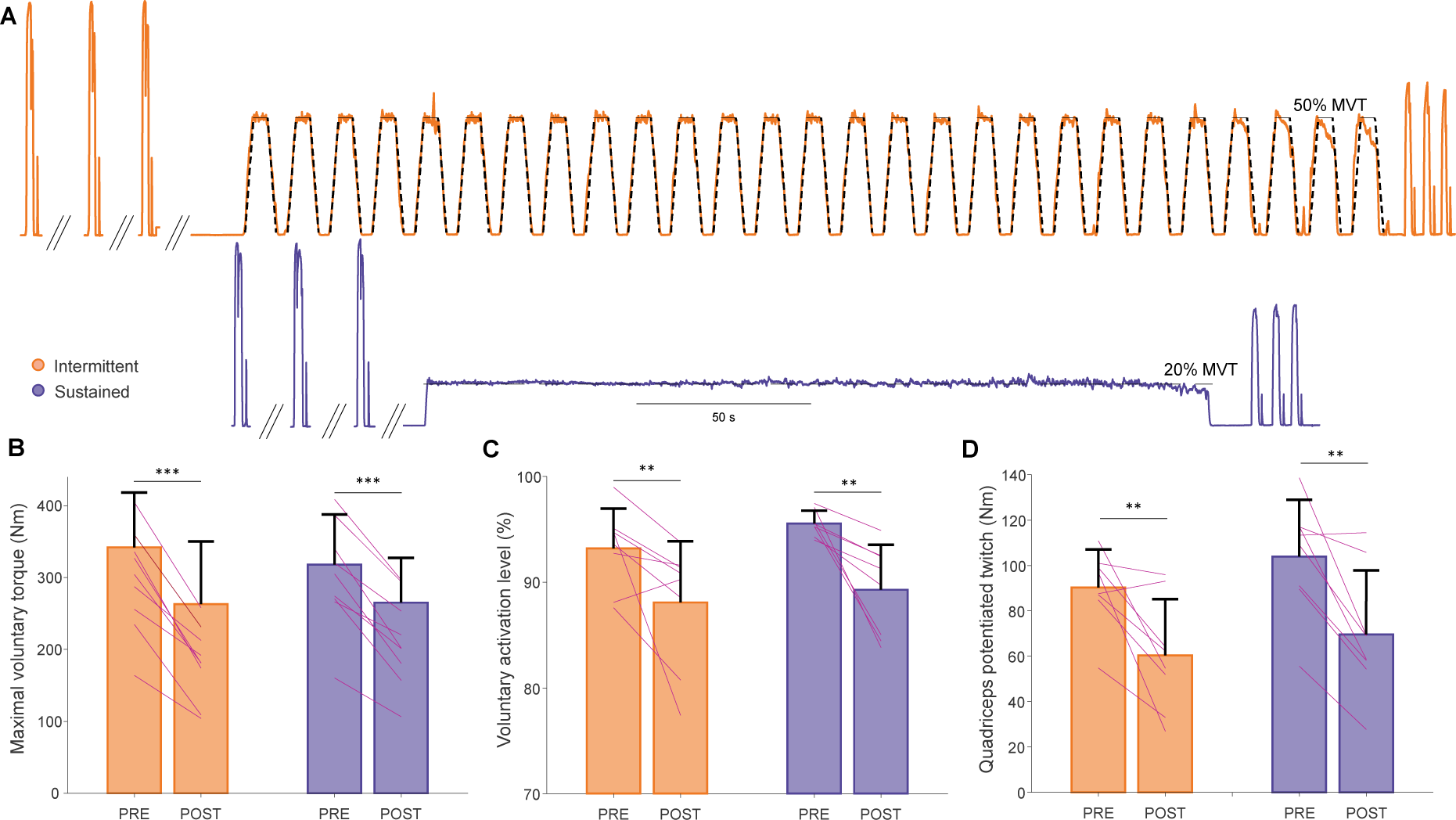
Task performance – torque, voluntary activation, and contractile properties. A: Example torque traces during the performance of the intermittent task (orange) at 50% of maximal voluntary torque (MVT) and the sustained task at 20% MVT (purple) to failure, with maximal voluntary contractions performed before and after each task and quadriceps twitches evoked during and immediately after each maximal contraction using femoral nerve stimulation. B: Maximal voluntary torque (B), voluntary activation level (C), and quadriceps potentiated twitch (D) before (PRE) and after (POST) the performance of the two contraction tasks to failure. Bar graphs indicate the mean with error bars denoting standard deviation, with lines indicating individual participant data. ***p < 0.001, **p < 0.01 compared to the other time point.

#### Torque steadiness and global EMG activity

Coefficient of variation of torque was predicted by contraction time (F_9, 142.0_ = 18.0, p < 0.0001), with it being greater than baseline at task failure for the intermittent task (p = 0.0021). During the sustained task, coefficient of variation of torque increased above baseline at 80% of contraction time (p = 0.0136), followed by a continuing increase until task failure (p < 0.0001).

The normalised root mean square EMG amplitude was predicted by contraction time (F_9, 271.8_ = 56.5, p < 0.0001), task (F_1, 275.3_ = 1112.6, p < 0.0001), and muscle (F_1, 272.4_ = 9.8, p = 0.0019), contraction time by task interaction (F_9, 271.8_ = 3.4, p = 0.0006), and task by muscle interaction (F_1, 272.7_ = 40.3, p < 0.0001). During the intermittent task, global EMG activity was greater between 60-100% of contraction time in VL (p = 0.0016 – p < 0.0001), and between 50-100% of contraction time in VM compared to baseline (p = 0.0125 – p < 0.0001). During the sustained task, global EMG activity was greater between 80-100% of contraction time compared to baseline in both VL (p = 0.0301 – p < 0.0001) and VM (p = 0.0272 – p < 0.0001). Global EMG activity was greater during the intermittent compared to the relative time points during the sustained task in both VL (p ≤ 0.0118) and VM (p < 0.0001).

**Figure 4.**
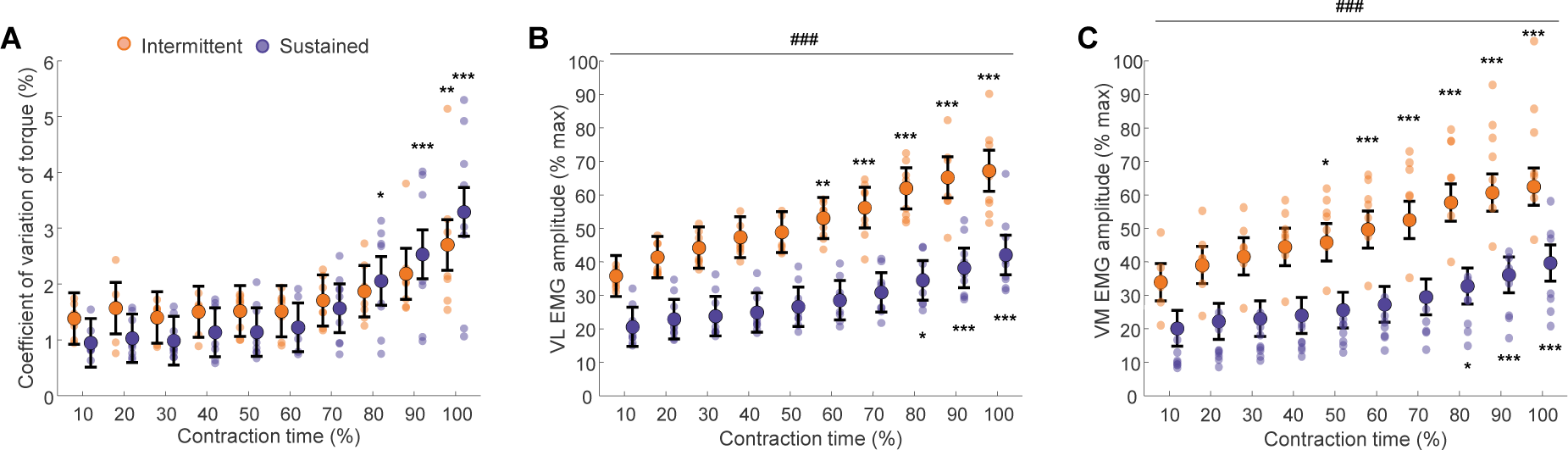
Torque steadiness and EMG amplitude changes during the performance of intermittent and sustained isometric task to failure. A, coefficient of variation of torque; B, C: vastus lateralis (VL, B) and medialis (VM, C) root mean square amplitude of electromyography (EMG) amplitude normalised to the EMG amplitude during the performance of maximal efforts before the start of the tasks. The bigger circles denote the estimated marginals means with the error bars denoting their 95% confidence intervals. The shaded, smaller circles represent individual participant scores (n = 9, 1 female). ***p < 0.001, **p < 0.01, *p < 0.05 compared to start of the task; ^###^p < 0.001 compared to the other task.

#### Motor unit identification

During the sustained task, 29 and 18 MUs were identified at the beginning of the task in VL and VM, respectively (4 ± 3 and 3 ± 2 per participant, respectively). With additionally recruited MUs throughout the task, 123 and 105 MUs were identified in VL and VM at the point of task failure (14 ± 9 and 13 ± 4 per participant, respectively). The discrepancy in the number of identified MUs between the beginning and end of the task highlights the previously reported difficulty in identifying MUs with smaller potentials in the presence of MUs with bigger potentials (Francic & Holobar, 2021). During the intermittent task, 49 and 48 early recruited MUs were identified in VL and VM (6 ± 7 and 7 ± 4 per participant), respectively, with the number of identified MUs increasing throughout the task to 80 and 81 MUs at task failure (10 ± 7 and 12 ± 6 per participant), respectively.

#### Motor unit discharge modulation during intermittent and sustained tasks

Discharge rate was modulated during both tasks. Examples of a few selected motor units in two participants during the isometric and intermittent task can be appreciated in Figure 5A and 5B, respectively. As can be noted, the later recruited MUs exhibited similar time-dependent changes in discharge rate to early recruited MUs.

**Figure 5.**
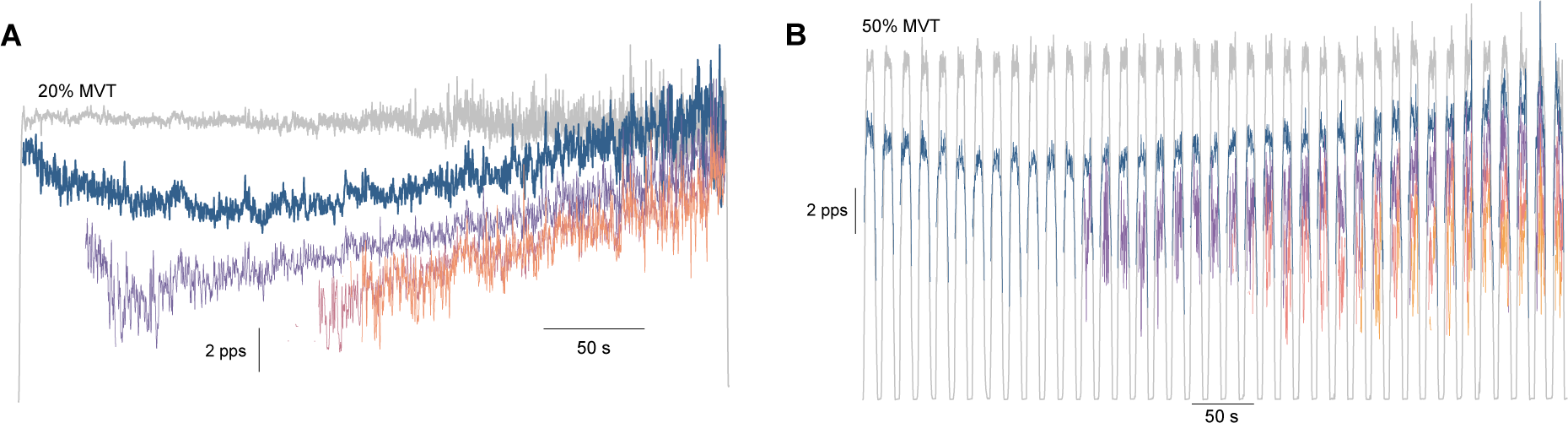
Example of motor unit discharge rate modulation during the tasks. Examples of smoothed discharge rates (400 ms Hann window) of a selected number of motor units (coloured) in vastus lateralis (VL) during the performance of a sustained (at 20% of maximal voluntary torque, MVT; A) and an intermittent (at 50% MVT; B) isometric knee extension tasks to failure. Note that the later recruited motor units seemingly exhibited mirroring changes in discharge rate to early recruited units.

Discharge rate was predicted by contraction time (F_9,3065.1_ = 141.4, p < 0.0001), task (F_1, 3071.6_ = 2273.5, p < 0.0001), muscle (F_1, 3067.5_ = 13.1, p = 0.0003), and recruitment time (F_1, 3072.4_ = 1027.7, p < 0.0001). There was also a significant interaction between contraction time and task (F_9, 3065.1_ = 62.2, p < 0.0001), and task and muscle (F_1, 3066.7_ = 16.2, p < 0.0001). During the sustained task, discharge rate initially decreased and was lower than at the beginning of the task at 40 (VL, p = 0.0231; VM, p = 0.0064), 50 (VL, p = 0.0040; VM, p = 0.0058), and 60% (VL, p = 0.0181; VM, p = 0.0299) of contraction time, after which it returned to baseline (p ≥ 0.0823; Figure 6A). During the intermittent task, discharge rate was similar to baseline up to 50 and 40% of contraction time in VL and VM, respectively (p ≥ 0.4264), at which point it increased above baseline (VL, p < 0.0001; VM, p = 0.0026), and continued to increase until task failure (p < 0.0001 for both VL and VM compared to baseline; Figure 6B).

**Figure 6.**
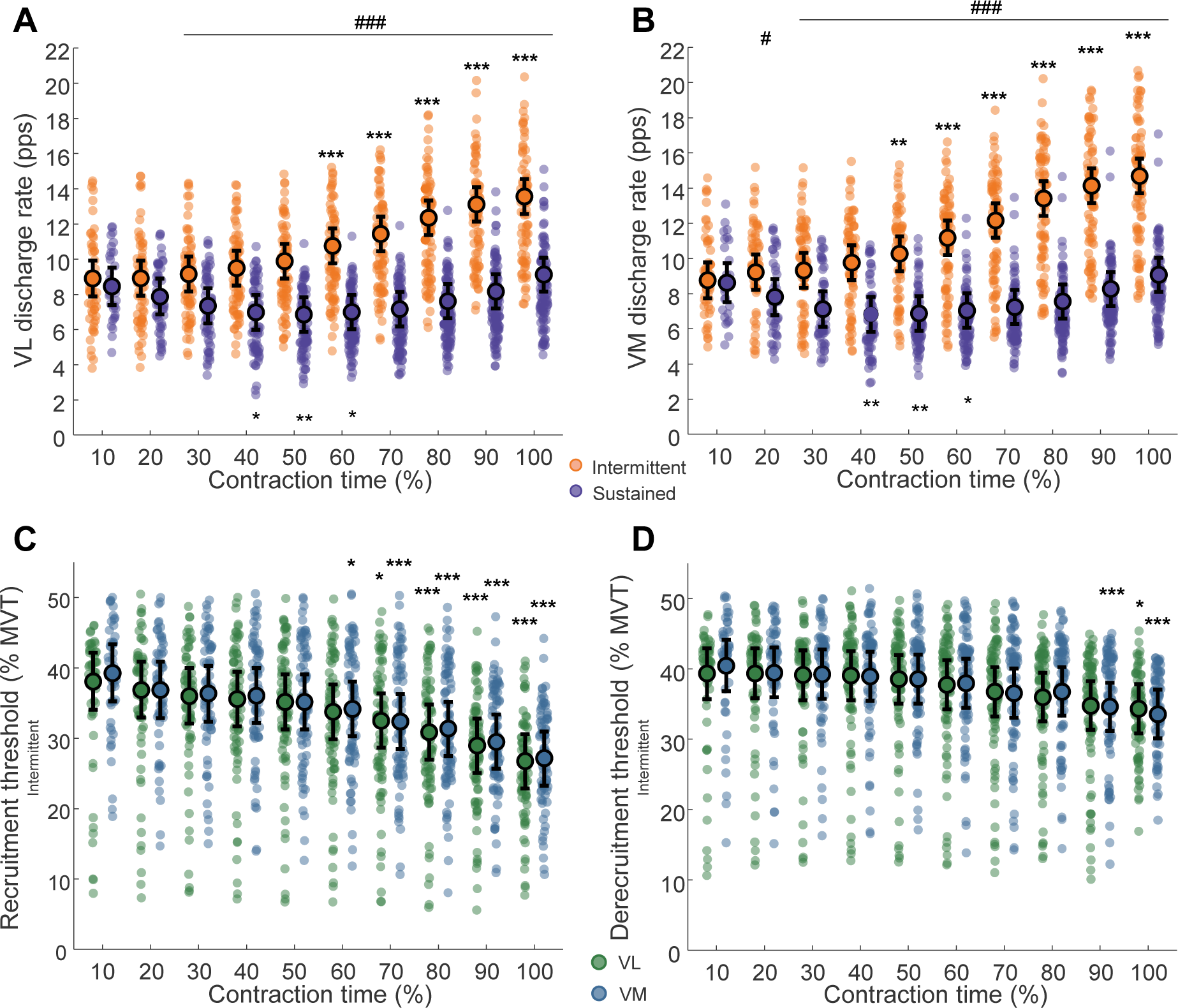
Motor unit discharge rate and (de)recruitment thresholds. A, B: Vastus lateralis (VL, A) and medialis (VM, B) motor unit discharge rates during the performance of intermittent (orange) and sustained (purple) isometric tasks to failure; C, D: motor unit recruitment (C) and derecruitment (D) thresholds during the performance of the intermittent task to failure. The bigger circles denote the estimated marginals means with the error bars denoting their 95% confidence intervals. The shaded, smaller circles represent individual motor units from all participants (n = 9, 1 female). ***p < 0.001, **p < 0.01, *p < 0.05 compared to start of the task; ^###^p < 0.001, ^#^p < 0.05 compared to the other task.

During the intermittent task, both recruitment (F_9, 1435.1_ = 31.0, p < 0.0001) and derecruitment (F_9, 1435.1_ = 11.4, p < 0.0001) thresholds of the identified MUs were predicted by contraction time. Recruitment threshold of the identified MUs was similar until 60 and 50% of contraction time in VL and VM, respectively, after which it decreased below baseline and continued to decrease until task failure (VL: p ≤ 0.0104; VM: p ≤ 0.0429; Figure 6C). Derecruitment threshold of the identified MUs was unchanged until 90 and 80% of contraction time in VL and VM, respectively, after which it decreased below baseline (VL: p = 0.0191; VM: p ≤ 0.0013; Figure 6D).

### Experiment 2

#### Time to task failure and decline in neuromuscular function

Time to task failure was 145 (26) and 72 (9) seconds for the 30 and 50% MVT sustained tasks (t_7_ = 9.9, p < 0.0001), respectively. The knee extensor MVT (F_1, 21_ = 170.2, p < 0.0001; Figure 7A), voluntary activation (F_1, 21_ = 10.1, p = 0.0046; Figure 7B), and quadriceps potentiated twitch torque (F_1, 21_ = 163.6, p < 0.0001; Figure 7C) decreased after both tasks, with no interactions between contraction time and task found for any of these variables (p ≥ 0.2578).

**Figure 7.**
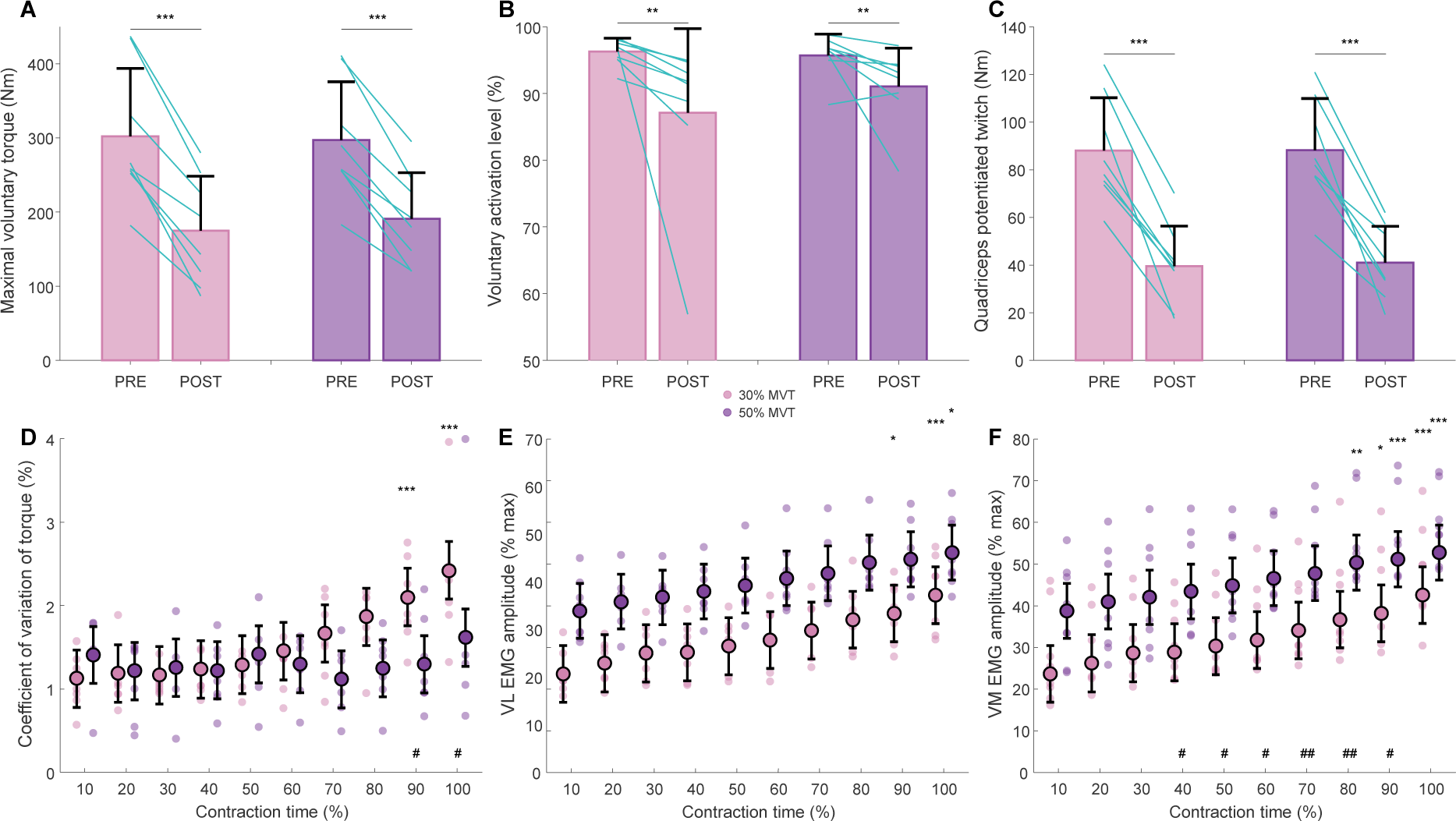
Torque, voluntary activation, contractile properties, and global EMG activity. Maximal voluntary torque (A), voluntary activation level (B), and quadriceps potentiated twitch (C) before (PRE) and after (POST) the performance of sustained contraction tasks at 30% and 50% of maximal voluntary torque (MVT) to failure. Bar graphs indicate the mean with error bars denoting standard deviation, and lines indicating individual participants’ data. ***p < 0.001, **p < 0.01 compared to the other time point. D, coefficient of variation of torque; E, F: vastus lateralis (VL, B) and medialis (VM, C) root mean square amplitude of electromyography (EMG) amplitude normalised to the EMG amplitude during the performance of maximal efforts before the sustained tasks to failure. The bigger circles denote the estimated marginals means with the error bars denoting their 95% confidence intervals. The shaded, smaller circles represent individual participants’ data (n = 8, 1 female). ***p < 0.001, **p < 0.01, *p < 0.05 compared to start of the task; ^###^p < 0.001, ^##^p < 0.01, ^#^p < 0.05 compared to the other task.

#### Torque steadiness and global EMG activity

Coefficient of variation of torque was influenced by contraction time, task, and their interaction. Post hoc testing showed that coefficient of variation of torque was greater compared to baseline at 90-100% of contraction time during the 30% MVT sustained task (p ≤ 0.0008), whereas it remained unchanged throughout the 50% MVT sustained task (p ≥ 0.9968; Figure 7D). The normalised root mean square EMG amplitude was predicted by contraction time (F_9, 263.0_ = 22.5, p < 0.0001), task (F_1, 263.3_ = 295.1, p < 0.0001), and muscle (F_1, 263.3_ = 11.2, p = 0.0009). During the 30% MVT sustained task, EMG amplitude was greater in the last 20% of contraction time compared to baseline in VL (p ≤ 0.0498) and VM (p ≤ 0.0151), respectively (Figure 7E). During the 50% MVT sustained task, EMG was greater than at baseline in the final 10 (p = 0.0317) and 30% (p ≤ 0.0020) of contraction time in VL and VM, respectively.

#### Motor unit identification

During the beginning of the 30% MVT sustained task, 41 and 34 (6 ± 8 and 4 ± 4 per participant) MUs were identified in VL and VM, respectively. The number of identified MUs increased to 79 and 55 (11 ± 12 and 7 ± 5 per participant) in VL and VM, respectively. During the 50% MVT sustained task, 46 and 44 MUs were identified at the beginning (6 ± 6 and 6 ± 5 per participant), and 53 and 70 MUs at the end of the task (7 ± 8 and 9 ± 8 per participant) in VL and VM, respectively.

#### Motor unit discharge modulation during sustained tasks

Discharge rate was predicted by contraction time (F_9,2222.1_ = 36.4, p < 0.0001), task (F_1, 2224.8_ = 487.5, p < 0.0001), muscle (F_1, 2224.3_ = 53.7, p < 0.0001), and recruitment time (F_1, 2227.2_ = 2850.2, p < 0.0001). There was also a significant interaction between contraction time and task (F_9, 2222.1_ = 2.8, p = 0.0026), and task and muscle (F_1, 2226.1_ = 47.7, p < 0.0001). Compared to baseline, VL discharge rate increased in the final 10% of the 30% MVT sustained task (p = 0.0036), whereas it was greater in the final 20% contraction time of the 50% MVT sustained task (p ≤ 0.0216; Figure 8A). Similarly, VM discharge rate was greater compared to baseline in the final 20 (p ≤ 0.0100) and 30% (p ≤ 0.0165) contraction time of the 30 and 50% MVT sustained tasks, respectively (Figure 8B).

**Figure 8.**
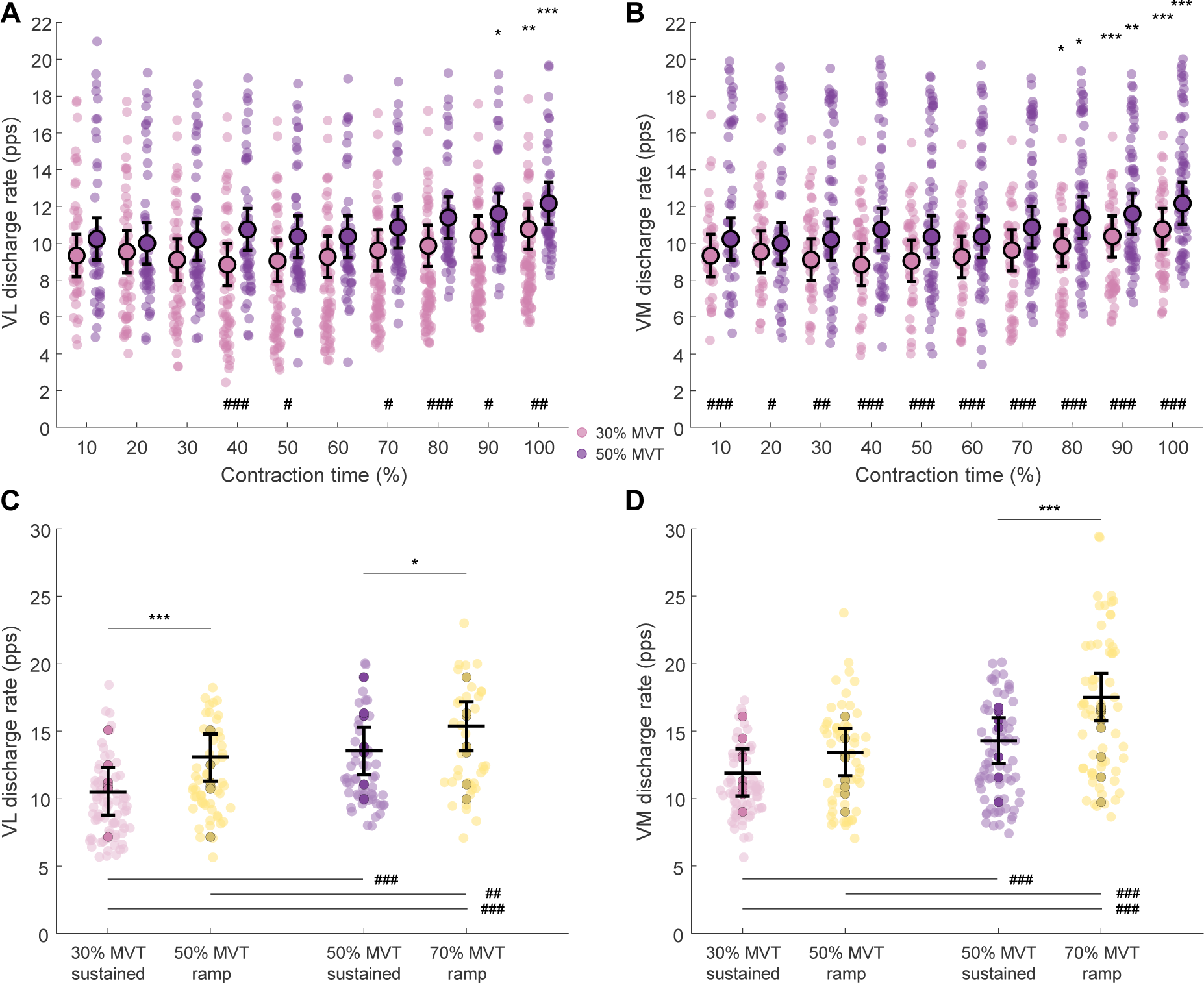
Motor unit discharge rate. A, B: Vastus lateralis (VL, A) and medialis (VM, B) motor unit discharge rates during the performance of sustained isometric task to failure performed at 30% and 50% maximal voluntary torque (MVT); The bigger circles denote the estimated marginals means with the error bars denoting their 95% confidence intervals. The shaded, smaller circles represent individual motor units from all participants (n = 8, 1 female). ***p < 0.001, **p < 0.01, *p < 0.05 compared to start of the task; ^###^p < 0.001, ^##^p < 0.001, ^#^p < 0.05 compared to the other task. C, D: VL (C) and VM (D) motor unit discharge rate during the final 10 seconds of the 30 and 50% MVT sustained task, compared to that during a trapezoidal ramp contraction (10% MVT/s increase/decrease, 10-second plateau) performed before the sustained tasks at a level 20% greater than the sustained task (i.e., 50% MVT for 30% MVT sustained task, and 70% MVT for 50% MVT sustained task). The coloured circles represent individual participant averages, the opaque circles denote values of individual motor units, whereas the horizontal black lines denote the estimated marginal means obtained from the linear mixed statistical modelling with the error bars denoting a 95% confidence interval. ***p < 0.001, **p < 0.01, *p < 0.05 relative to the other contraction, ^###^p < 0.001, ^##^p < 0.01, ^#^p < 0.01 relative to contractions related to the other task.

When comparing contractions before the tasks at a level 20% MVT greater than the sustained tasks (i.e. 50% MVT for the 30% MVT task and 70% MVT for the 50% MVT task) with the final 10 seconds of the tasks to failure, discharge rate was predicted by contraction (sustained vs. trapezoidal ramp; F_9,459.8_ = 76.0, p < 0.0001), task (F_1,460.0_ = 128.9, p < 0.0001), and muscle (F_1,461.8_ = 18.5, p < 0.0001). There was also an interaction between contraction, task, and muscle (F_1,459.6_ = 5.5, p = 0.0192). In VL, discharge rate at task failure was still greater during trapezoidal ramp contractions performed before the sustained tasks compared to that at task failure (30% MVT sustained task vs. 50% MVT trapezoidal ramp contraction: p < 0.0001; 50% MVT sustained task vs. 70% MVT trapezoidal ramp contraction: p = 0.0339; Figure 8C). In VM, this was still the case when discharge rate at task failure during 50% MVT sustained task was compared to that during the 70% MVT trapezoidal ramp contraction (p < 0.0001), whereas this was not the case when discharge rate during 50% MVT trapezoidal ramp contraction was compared to that at task failure during the 30% MVT sustained task (p = 0.0705; Figure 8D).

## DISCUSSION

This study examined task-specificity of MU discharge behaviour adjustments in VL and VM during prolonged isometric knee extension tasks to failure. Two independent experiments were performed. Experiment 1 compared MU discharge rate modulation pattern during a sustained and intermittent contraction task at 20 and 50% MVT, respectively, and Experiment 2 compared MU discharge rate adjustments during sustained contraction tasks at 30 and 50% MVT. In all conditions, we showed evidence of a biphasic pattern of MU discharge rate modulation. During the 20% MVT sustained contraction task, MU discharge rate initially decreased, which was followed by an increase such that MU discharge rate at task failure was similar to baseline. Conversely, no relative decrease in MU discharge rate was observed for the other contraction tasks (30% MVT sustained, and 50% MVT sustained and intermittent); rather, MU discharge rate remained similar for the first part of the task, followed by an increase towards task failure, with the greatest rate of increase observed during the intermittent task. Our results thus suggest that discharge rate modulation patterns during isometric muscle contractions to failure are task- and intensity-dependent.

### Differences in the initial motor unit discharge rate adjustments

The key difference between the intermittent and sustained tasks in the Experiment 1 was the decrease in MU discharge rate in the first half of the task. Specifically, a reduction in MU discharge rate was observed in the 20% MVT sustained task, but not during the 50% MVT intermittent task. However, when comparing MU discharge behaviour during sustained contractions at 30 and 50% MVT, there was no decrease in MU discharge rate during the initial period of any of the two tasks.

The observed MU discharge rate behaviour during the 20% MVT sustained task is similar to Martinez-Valdes and colleagues (2020) who also observed a decrease (reaching significance at ∼40% contraction time) followed by an increase in VL and VM MU discharge behaviour during a 30% MVT sustained task. However, we could not replicate these results when performing a 30% sustained MVT task. The discrepant results may be, at least partially, attributable to subtle differences in approaches to analysis. These include differences in MU tracking methodology, and the inclusion/exclusion of later recruited MUs; for example, in our study we included later recruited MUs in the analysis, considering recruitment time as a covariate, whereas Martinez-Valdes and colleagues considered only MUs recruited from the beginning of the task (Martinez-Valdes *et al*., 2020a). Since MUs recruited later in the task, particularly those recruited beyond a point of general increase in MU discharge rate (see Figure 5A & 5B for example), are less likely to exhibit initial decline in MU discharge, this phenomenon might be less likely to be statistically detected.

The decrease in discharge rate in the first part of the 20% MVT sustained task is unlikely to have been caused by the inhibitory afferent feedback via group III/IV muscle afferents as this effect would have presumably been greater during higher intensity tasks that will have induced greater ischaemia and thus likely have a greater intramuscular accumulation of noxious metabolites. The observed behaviour is similar to the reduced responsiveness of motoneurons to constant synaptic input as previously shown in reduced animal preparations (i.e., late spike frequency adaptation; Kernell & Monster, 1982; Spielmann et al., 1993; Sawczuk et al., 1995; Button et al., 2007) and indirectly in humans (Finn *et al*., 2018; Brownstein *et al*., 2021). However, spike frequency adaptation seems a less likely explanation for the observed reduction in MU discharge rate in the initial part of the 20% MVT sustained task since such an effect has been shown to be more pronounced in higher threshold motoneurons (Spielmann *et al*., 1993; Button *et al*., 2007). This is inconsistent with our results showing the lack of relative decline in MU discharge rates at higher contraction levels (30 and 50% MVT) and the fact our data likely reflect the behaviour of mainly higher threshold MUs due to decomposition bias (Francic & Holobar, 2021; see also Figure 6C & 6D). Importantly however, evidence for late spike frequency adaptation largely stems from studies in reduced animal preparations in the absence of neuromodulatory inputs. Indeed, increasing monoaminergic input that facilitates persistent inward currents (PICs) on motoneuron dendrites has been shown capable of diminishing the effects of late spike frequency adaptation (Hornby *et al*., 2002; Brownstone *et al*., 2011). Considering the likely greater PIC magnitude with greater contraction intensity (Škarabot *et al*., 2023b), we postulate that PIC magnitude is insufficient at 20% MVT to prevent the effects of late spike frequency adaptation. An alternative explanation for the lack of observed decreases in MU discharge rate during tasks performed at higher contraction intensities is purely methodological, grounded in normalising contraction time in 10% intervals to allow comparison between participants, which inevitably acts as a smoothing function, making smaller or shorter/shallower decreases in MU discharge rates less likely to be detectable.

The more likely mechanism for the observed reduction in MU discharge rate in the initial part of the 20% MVT sustained task relates to MU twitch potentiation. Assuming an increase in MU twitch amplitude and area with initial repetitive depolarisation (Bagust *et al*., 1974; Burke *et al*., 1976), a constant torque output could be maintained despite decreased MU discharge rate. Indeed, the time course of twitch potentiation has been demonstrated to coincide with the decrease in MU discharge rate in the beginning of the submaximal isometric task to failure (Martinez-Valdes *et al*., 2020a). Eventually, MU twitch amplitude has been shown to decline (Martinez-Valdes *et al*., 2020a), likely due to the accumulation of metabolites (McKenna *et al*., 2008) that render muscle contractile capacity reduced, necessitating greater synaptic input to motoneurons to maintain the same torque output resulting in an increase in MU discharge rate until task failure. Whilst evidence in cats suggests that the extent of MU twitch potentiation is similar between lower and higher threshold MUs (Bagust *et al*., 1974), we speculate that the shorter time course of potentiation (during the 30% and 50% MVT sustained tasks) and interruption of repetitive activation (during the 50% MVT intermittent task) lead to a significantly smaller or less detectable decreases in MU discharge rate in the initial part of the task at higher intensity (note a representative example of an early recruited MU in Figure 5B that demonstrates a marginal trend of decrease in discharge rate). Overall, these results suggest that MU discharge rate behaviour in the first portion of the task is more related to intensity-, rather than the task.

### Inability to increase motor unit discharge rate coincides with task failure

During all tasks, MU discharge rate progressively increased across the second half of the task. During the sustained contraction at 20% MVT, the increase began at a below-baseline level (due to the initial decrease) and did not increase above baseline at task failure. During the sustained contractions at 30 and 50% MVT and the intermittent contractions at 50% MVT, MU discharge rate increased above baseline, with the degree of the increase being dependent on the contraction intensity, being lesser in 30% compared to 50% MVT sustained tasks. increase in MU discharge rate in the latter parts of the task, however, appeared to be the greatest during the 50% MVT intermittent task to failure, where the increase above baseline was detected at the earliest point in the task (∼60% contraction time) and kept increasing until task failure. These observations suggest that the relative changes in MU discharge behaviour are both intensity- and task-dependent.

The increase in MU discharge rate with proximity to task failure likely reflects an increase in the strength of the common excitatory synaptic input (Castronovo *et al*., 2015), which is necessary to maintain constant torque output, likely coinciding with the decrease in MU twitch potentiation (Martinez-Valdes *et al*., 2020a). There are several lines of observation to support this interpretation. First, with increased strength of the common synaptic input, torque variability tends to increase (Castronovo *et al*., 2015); indeed, we demonstrated greater coefficient of variation of torque towards the end of the tasks. Second, the greater strength of common synaptic input likely reduces the membrane potential needed for depolarisation, reducing the threshold of already recruited MUs, and resulting in recruitment of additional MUs, both of which were demonstrated in our data. MU recruitment thresholds decreased with contraction time during the intermittent task, contrasting the changes in MU discharge rate; recruitment remained stable during the first half of the task, and then decreased below baseline (coinciding with the start of MU discharge rate increase) and continue to decrease until task failure. This is consistent with previous findings in an intermittent task (Carpentier *et al*., 2001; Farina *et al*., 2009) and suggests that, with repetitive motoneuron activation, MU recruitment range progressively compresses. The reduction in recruitment threshold coupled with the increase in MU discharge rate likely represents a neural strategy which serves to maintain constant torque output in the presence of impaired contractile function. Along similar lines, derecruitment thresholds were also reduced, though later in the intermittent task, indicative of prolonged MU discharge as a result of greater common synaptic input. Finally, though several later recruited MUs exhibited a constant increase in MU discharge rate, this was dependent on their recruitment time within the task. For example, if additional MUs were recruited relatively early in the task, their behaviour was consistent with already recruited MUs (note the example in Figure 5A, whereby an additionally recruited MU early in the task initially exhibits a decrease in discharge rate followed by an increase, akin to the MUs recruited in the beginning of the task). These observations support the notion that the common synaptic input strength is modulated throughout the task to allow constant torque output.

During submaximal sustained contractions at 30 and 50% MVT, MU discharge rate increased above baseline in the final 10 and 20% contraction time, respectively. However, MU discharge rate at task failure did not reach the rates exhibited during a short, higher-level (+20% MVT) contraction performed before contractions to failure, in agreement with previous findings (Martinez-Valdes *et al*., 2020a). Similarly, global EMG amplitude at task failure, though likely more reflective of MU recruitment and peripheral alterations (Del Vecchio *et al*., 2017; Martinez-Valdes *et al*., 2018), also did not reach EMG amplitude recorded during MVT, consistent with prior work (Fuglevand *et al*., 1993). These findings suggest that the increase in common synaptic input is ultimately insufficient to allow depolarisation and continued increases in MU discharge rate, coinciding with task failure. The relative increase in MU discharge rate at task failure seemed to be greater during the intermittent task, and during tasks performed at greater contraction intensity, suggesting different mechanistic interaction depending on task and intensity leading to the observed MU discharge behaviour. Evidence from reduced animal preparations shows that an intense or prolonged release of serotonin may cause a spillover onto the extrasynaptic receptors on the axon initial segment and thus reduce motoneuron activity (Cotel *et al*., 2013; Perrier *et al*., 2018). However, whilst human studies have indicated the potential for ingestion of serotonin reuptake inhibitor to shorten time to task failure (Kavanagh *et al*., 2019), this seems only to be the case for maximal, rather than submaximal sustained contractions (Henderson *et al*., 2022), making it an unlikely mechanism to explain the observed differences in relative increase in MU discharge rate between tasks and contraction intensities in our study.

The task-dependency of MU discharge behaviour is perhaps unsurprising; the greatest increase in MU discharge rate during the intermittent task is likely due to lesser levels of ischaemia, with the post-contraction hyperaemic response restoring muscle perfusion and clearing accumulated metabolites (Adreani & Kaufman, 1998; De Ruiter *et al*., 2007), thus reducing the inhibitory inputs of group III/IV afferents to motoneurons (Martinez-Valdes *et al*., 2020b) and the motor cortex (Kennedy *et al*., 2016) compared to that during the sustained contractions. Similarly, H-reflex has been shown to reduce during sustained compared to intermittent contractions (Duchateau & Hainaut, 1993), indicating reduced Ia afferent input to motoneurons, likely further contributing to smaller increases in MU discharge rate during sustained contractions. Though not observed in our study, the smaller increases in MU discharge rate during lower intensity sustained contractions are consistent with the notion that reductions in voluntary activation are typically greater during lower relative torque outputs (Burnley *et al*., 2012), indicating a greater impairment in the central drive. During submaximal sustained contractions, corticospinal excitability and inhibition have been shown to progressively increase (Søgaard *et al*., 2006; Lévénez *et al*., 2008; Klass *et al*., 2008), whereas such changes appear intensity-dependent during intermittent contractions, with corticospinal excitability increased and decreased during contraction above and below critical torque, respectively (Ansdell *et al*., 2019a). Future studies should consider employing such measures concurrently with HDsEMG to attempt to explain the apparent contraction intensity-dependent alterations in MU discharge rate in isometric tasks to failure.

### Further considerations

Previous studies investigating MU discharge behaviour during prolonged contractions have produced mixed findings. Although the consensus sometimes appears to suggest a progressive decline in MU discharge rate should be expected with sustained contractions (Enoka & Duchateau, 2008), comparisons must be interpreted through the lens of methodological differences, the nature of the task, the motor pool investigated, and possibly the recruitment threshold of the identified MUs, among others. The studies that did not perform contraction tasks to failure seem less likely to be comparable; the progressive decrease of MU discharge rate in those studies (Mottram *et al*., 2005; Riley *et al*., 2008; Pascoe *et al*., 2014) might reflect task cessation before the MU twitch potentiation had reached its peak, not necessitating the need for an increase in synaptic input to motoneurons to maintain the constant torque output. Our demonstration of a biphasic behaviour in MU discharge, with an increase towards task failure seems to agree with a large number of studies that performed tasks to failure (Bigland-Ritchie *et al*., 1986; Garland *et al*., 1997; Griffin *et al*., 2000; Kuchinad *et al*., 2004; Adam & De Luca, 2005; Mettler & Griffin, 2016), though not with some others (Garland *et al*., 1994; Carpentier *et al*., 2001; McManus *et al*., 2016).

It is also worth considering methodological differences between studies, with many prior studies utilising intramuscular EMG and template matching algorithms that carry a higher risk of erroneous MU identification during prolonged contraction where MU action potential waveforms are likely to change due to modulation in conduction velocity (Farina *et al*., 2009). In this study, we utilised an approach of estimating and applying MU filters in successive, short windows, which has shown to be a robust approach when attempting to accommodate small changes in MU waveforms (Kramberger & Holobar, 2021; Francic & Holobar, 2021; Škarabot *et al*., 2023a). From this perspective, our results broadly agree with a study that utilised multichannel surface EMG decomposition, and a time-domain MU tracking approach (Martinez-Valdes *et al*., 2020a). Furthermore, the number of concurrently active MUs identified with intramuscular EMG is limited, which may underestimate the modulation in MU discharge rate at the population level (Enoka, 2019). Finally, many of the prior studies pooled the identified MU discharges, violating the principle of co-dependence of observations (Tenan *et al*., 2014). Conversely, we used linear mixed statistical modelling, accounting for nesting of observations within individual participant (Yu *et al*., 2022; Wilkinson *et al*., 2023). Finally, in most of the studies showing a progressive decrease in MU discharge rate to task failure, an intrinsic hand muscle was used (Carpentier *et al*., 2001; McManus *et al*., 2016). Considering previous suggestions that the behaviour of MU discharge rate adjustments with repetitive motoneuron activation might depend on the upper limit of recruitment of the motor pool (Enoka & Stuart, 1992), we allow for the possibility that the demonstrated intensity- and task-dependent MU behaviour in our study might be constrained to the vastii motor pools, or at least motor pools with similar upper limits of recruitment. It remains unclear whether the results can be replicated in a similar design using an intrinsic hand muscle with a comparatively lower upper limit of MU recruitment (Kukulka & Clamann, 1981).

It is important to note that owing to decomposition bias, the majority of the identified MUs were of higher threshold, and thus we cannot exclude the possibility that the observed behaviour is related to MU recruitment threshold, rather than intensity of the tasks per se. For example, during intermittent contractions where recruitment thresholds could be reliably determined, most of the identified MUs in the beginning of the task had recruitment thresholds >35% MVT on average. It is conceivable that MU behaviour exhibited during repetitive activation of motoneurons is nonhomogeneous across the motor pool, with previous reports showing a decrease in discharge rate of early recruited, but an increase in later recruited (higher threshold) MUs (Carpentier *et al*., 2001). We provide some evidence of this insofar as we observed later recruited MUs (that will have higher thresholds) to have a tendency to increase discharge rate. Future studies identifying a pool of MUs with a sufficient range of recruitment thresholds, perhaps by combining intramuscular and surface EMG recordings, or increasing the density and/or area of surface recordings (Caillet *et al*., 2023), might be able to answer this question more directly.

## Conclusion

We showed that during isometric contractions to failure, vastus lateralis and medialis MU discharge rate alterations exhibit distinct, biphasic patterns depending on contraction task and intensity. During a low intensity (20% MVT) contraction, MU discharge rate initially decreased, before increasing to baseline levels. During higher intensity contractions, either sustained at 30 and 50% MVT, or intermittent at 50% MVT, we showed that MU discharge rate remained stable, after which it continually increases until task failure, though the relative increase depends on contraction intensity, with the greatest increase demonstrated by the intermittent task. In all tasks, modulations in MU discharge rate pattern were also accompanied by recruitment of additional MUs. The observed behaviour is likely related to changes in the strength of the common excitatory synaptic input that is modulated in response to MU twitch potentiation and attenuation throughout the isometric task to failure. Notably, the increased MU discharge rate at task failure for the 30% and 50% MVT sustained tasks was still lower than during short trapezoidal contractions performed at 50% and 70% MVT, respectively, suggesting insufficient strength of the excitatory synaptic input to motoneurons to allow further discharge rate increases and prolongation of task. Overall, the data presented herein provides strong evidence that MU discharge rate modulation during isometric contractions to failure is both intensity- and task-dependent.

## Author contributions

TV, PA, CGB, AH, and JŠ conceived and designed research; TV and JŠ performed experiments; TV, PS, AH, and JŠ analysed data; TV and JŠ drafted manuscript; all authors edited and revised manuscript and approved the final version.

